# Dual RNA-seq identifies proteins and pathways modulated during *Clostridioides difficile* colonisation

**DOI:** 10.1101/2023.01.30.526182

**Authors:** Lucy R. Frost, Richard Stark, Blessing Anonye, Ludmila R. P. Ferreira, Meera Unnikrishnan

**Affiliations:** Division of Biomedical Sciences, Warwick Medical School, University of Warwick, Coventry CV47AL, United Kingdom; Bioinformatics Research Technology Platform, University of Warwick, Coventry CV47AL, United Kingdom; RNA Systems Biology Laboratory (RSBL). Department of Morphology, Institute of Biological Sciences, Federal University of Minas Gerais, Belo Horizonte, MG 31270-910, Brazil

**Keywords:** *Clostridioides difficile* infection, *in vitro* gut model, dual RNAseq, proline-proline endopeptidase

## Abstract

The gastrointestinal pathogen, *Clostridioides difficile*, is the most common cause of hospital-acquired diarrhoea. Bacterial interactions with the gut mucosa are crucial for colonisation and establishment of *C. difficile* infection, however, key infection events like bacterial attachment and gut penetration are still poorly defined. To better understand the initial events that occur when this anaerobic pathogen interacts with human gut epithelial cells, we employed a dual RNA-sequencing approach to study the bacterial and host transcriptomic profiles during *C. difficile* infection in a dual-environment *in vitro* human gut model. Temporal changes in gene expression during infection were studied in the bacterium and the host cells over the course of 3-24 hours. While there were several common differentially expressed bacterial genes across the different times after infection, mammalian transcriptional profiles were quite distinct with little overlap. Interestingly, an induction of colonic receptors for *C. difficile* toxins was observed, along with the expression downregulation of genes encoding immune response markers. Several cell wall associated proteins were downregulated in *C. difficile* when in association with host cells, including *slpA*, which encodes the main S-layer protein. Gene function and pathway enrichment analyses revealed a potential modulation of the purine/pyrimidine synthesis pathways both in the mammalian and the bacterial cells. We observed that proline-proline endopeptidase, a secreted metalloprotease responsible for cell surface protein cleavage, is downregulated during infection, and a mutant lacking this enzyme demonstrated enhanced adhesion to epithelial cells during infection. This study provides new insight into the host and bacterial pathways based on gene expression modulation during the initial contact of *C. difficile* with gut cells.

**Importance:** The initial interactions between the colonic epithelium and the bacterium are likely critical in the establishment of *Clostridioides difficile* infection, one of the major causes of hospital acquired diarrhoea worldwide. Molecular interactions between *C. difficile* and human gut cells have not been well defined mainly due to the technical challenges of studying cellular host-pathogen interactions with this anaerobe. Here we have examined transcriptional changes occurring in the pathogen and host cells during the initial 24 hours of infection. Our data indicate several changes in metabolic pathways and virulence-associated factors during the initial bacterium-host cell contact and early stages of infection. We describe canonical pathways enriched based on the expression profiles of a dual RNAseq in the host and the bacterium, and functions of bacterial factors modulated during infection. This study provides insight into the early infection process at a molecular level.

## Introduction

*Clostridioides difficile* is a Gram-positive, anaerobic, spore forming bacterium and one of the most frequently reported nosocomial intestinal pathogens. Clinical manifestations of *C. difficile* infection (CDI) can range from mild diarrhoea to pseudomembranous colitis and toxic megacolon, which are often life threatening (1). In the UK, although the number of cases of CDI reported have been declining in recent years, there were 14,249 cases reported in the UK between 2021/2022 (2). However, in the USA, the number of cases is currently very high, with approximately half a million cases being reported each year, and an estimated 14,000 deaths (3, 4). High recurrence rates of CDI have placed a huge cost burden on healthcare systems worldwide (5).

*C. difficile* produces highly resistant spores which germinate in the presence of bile salts in the gastrointestinal tract (6). Vegetative cells colonise the gut mucosa when the normal gut microbiota is disrupted, usually after antibiotic therapy. Following successful colonisation of the gut epithelium, *C. difficile* replicates and secretes toxins: the enterotoxin TcdA, the cytotoxin TcdB, as well as an ADP-ribosylating toxin (CDT) in ribotype 027 strains (7). These toxins are primarily responsible for epithelial barrier disruption, tissue damage and fluid accumulation during infection (8). In addition to toxins, several other bacterial factors have been identified which have exhibited adhesive properties to host extracellular matrix (ECM) components, epithelial cell layers or gastrointestinal tissues. These colonisation factors include surface layer proteins (SLPs), cell wall proteins (CWPs), flagella, pili, ECM-binding proteins, and secreted enzymes like the proline-proline endopeptidase (PPEP-1) (9-14). As seen with other pathogens, *C. difficile* is recognised by toll-like receptors (TLRs) including TLR-5 and TLR-2 on the host cell-surface, initiating innate immune responses against the bacterium (15-17). *C. difficile* infection is also accompanied by an increase in abundance of several proinflammatory cytokines, including interleukins (IL): IL-8, IL-1β, IL-23, IL-33 and tumor necrosis factor alpha (TNF-α) and chemokines, such as C-C motif chemokine ligand 5 (CCL-5) and CCL-2 (18-23). Early recruitment of immune cells, such as neutrophils, eosinophils, and interferon gamma (IFN-γ)-producing type 1 innate lymphoid cells, IL-17 producing ψο T cells and IL-33 (19, 24, 25) have also been associated with protection against *C. difficile* infection. Additionally, this bacterium can persist within the gut causing recurrent infections, with sporulation and biofilm formation being key processes contributing to persistence (6, 26).

To understand host pathways activated during infection, host transcriptomic responses have been studied in response to toxins A and B in murine models (27, 28). Studies have also examined the whole tissue transcriptional responses to vegetative *C. difficile* in a murine infection model, where IL-33 was shown to be upregulated during infection (19). Other studies have examined the bacterial transcriptional responses during infection in porcine and murine models, identifying several known virulence-associated factors, and revealing new colonisation factors (29, 30). Detailed studies on human intestinal cells have not been possible due to a paucity of *in vitro* models allowing for coculture of the anaerobic *C. difficile* with oxygen-requiring epithelial cells.

In this study, we investigated the global host and bacterial transcriptomic responses during early *C. difficile* infection, employing a dual environment (anaerobic and aerobic) *in vitro* human gut model (31) and dual RNA-seq. We describe here the differential expression of several genes from host and bacterial cells during the initial 24 h of *C. difficile* infection. Computational analyses of transcriptomic data identified the enriched canonical pathways at different timepoints after infection, including one related to purine/ pyrimidine synthesis, which was enriched in both *C. difficile* and host cells. We observed a downregulation of a proline-proline endopeptidase, a secreted metalloprotease responsible for cleavage of cell surface proteins and further demonstrated that it modulates bacterial adhesion to epithelial cells during infection.

## Results

### Dual RNA-seq of *C. difficile*-infected gut epithelial cells in an *in vitro* gut model

We have previously optimised an *in vitro* gut infection model where polarised epithelial layers comprising Caco-2 and HT29-MTX cells (9:1) were infected by *C. difficile* within a vertical diffusion chamber (VDC) system, which enables coculturing in anaerobic and aerobic environments (31). To study transcriptional responses in bacterial and epithelial cells simultaneously, the intestinal epithelial cell layers were infected with *C. difficile* in the VDC at a multiplicity of infection (MOI) of 10 for 3 h. Extracellular bacteria were washed off, and the cells with adherent bacteria were further incubated until 6, 12 and 24 h post infection (Fig. 1). To generate host uninfected control samples, cell layers were incubated in the VDC system for the required times without the addition of *C. difficile*. As a control for the bacterial samples, *C. difficile* culture was grown to log-phase in pre-reduced DMEM-10, the same medium used for the infections. Total RNA was extracted from all the infected samples and uninfected controls from 3, 6, 12 and 24 h incubation (Fig. 1) and DNA libraries were prepared and sequenced as described in Methods. Sequencing generated 49-64 million reads for the infected samples. While, for all infected samples, over 95% of reads align to a concatenated reference containing both the human (GRCh38) and *C. difficile* R20291 genomes, the read mapping proportion to the *C. difficile* genome alone was much lower, ranging from 0.06% to 2.06% (Table S1). Although bioanalyzer profiles of purified RNA indicated efficient removal of rRNA we noted that there was a high proportion of reads mapping to human rRNA upon sequencing.

**Figure 1.**
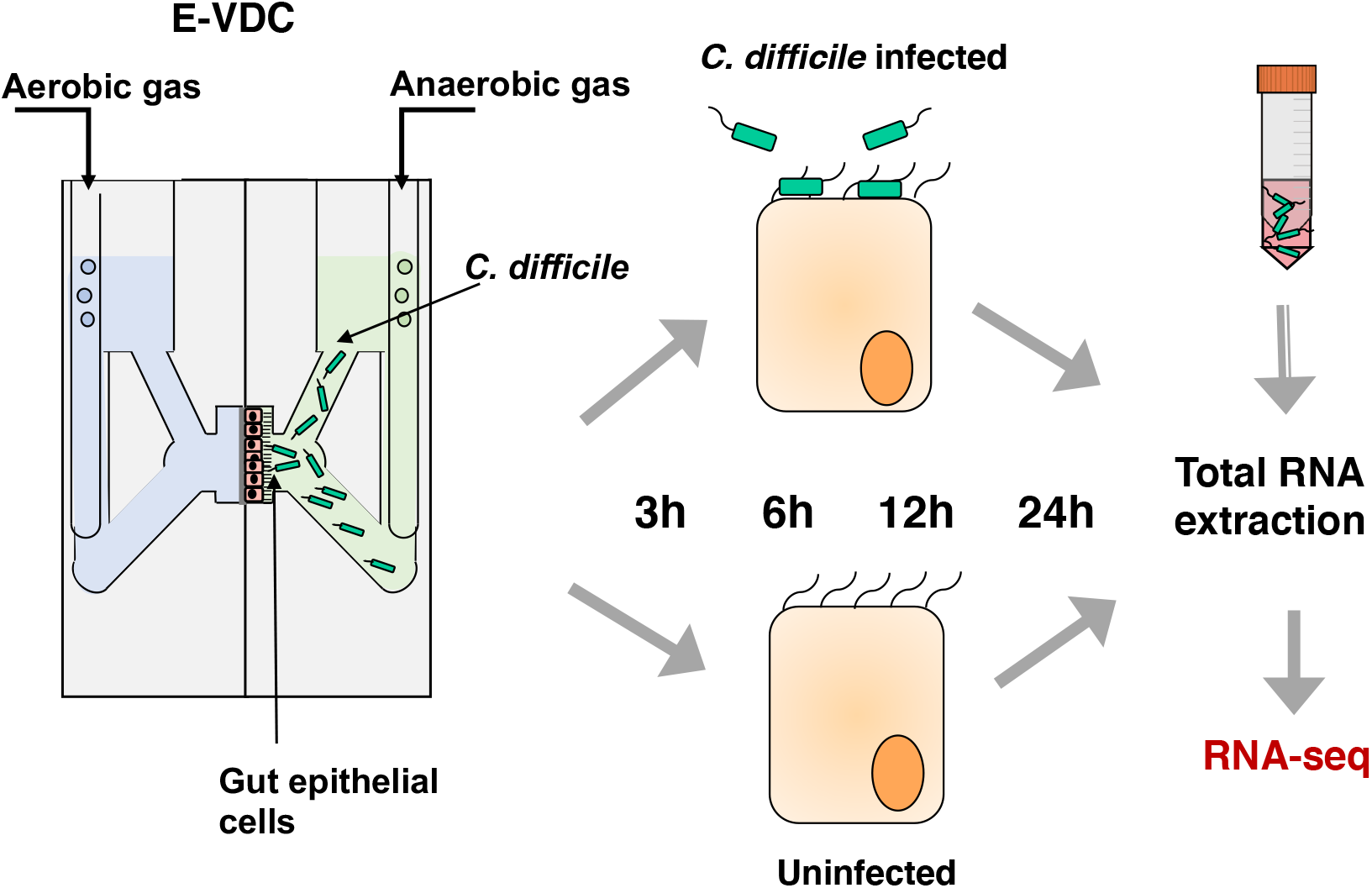
A schematic diagram of the dual RNA-seq experimental setup. An *in vitro* human gut model employing a vertical diffusion chamber (VDC) to facilitate *C. difficile* infection of intestinal epithelial cells. Total RNA was extracted from adhered bacteria and intestinal epithelial cells incubated in the system for 3, 6, 12 or 24 h, with or without *C. difficile*. A bacterial inoculum grown to logarithmic phase in pre-reduced DMEM-10 was used as an uninfected bacterial control.

PCA analysis for all samples with human cells displayed clustering of samples within treatment replicate groups at all timepoints (Fig S1A). A clear transcriptional shift was observed between uninfected controls and infected samples at all timepoints. We noted changes in the gene expression profiles over the time course in infected cell samples, but also in the uninfected human cell controls, albeit to a lesser degree. The uninfected 24 h controls were clustered more closely to the late-stage infected groups than the other uninfected control groups, suggesting a stress response in the intestinal epithelial cells incubated in this static system which is exacerbated over time. For the bacterial samples, PCA plots indicated that biological replicates of 3 and 6 h infected samples had large gene expression variations compared to the other replicates. For this reason, these outliers were completely removed from the analysis. Following their removal, all replicates within samples were clustered together and a clear transcriptional shift from the bacterial culture control was observed for all timepoints (Fig. S1B).

Differentially expressed genes (DEGs) were identified by comparing control to infected human samples for each time as described in Methods. A total of 205 and 196 human genes were significantly upregulated and downregulated respectively (adjusted p-value (p adj) < 0.05 and log_2_(fold change) greater than 1 or less than -1), with 121 upregulated, and 160 downregulated only at 3 h. No genes were significantly up- or downregulated at all timepoints (Fig 2A). In case of *C. difficile*, a total of 222 genes were significantly upregulated, of which 82 were upregulated at all timepoints, and 229 *C. difficile* genes were downregulated, of which 69 were significantly downregulated at all timepoints (Fig. 2B, Fig. S2). Thus, the mammalian transcriptional responses induced at each time point during infection appears to be distinct with little overlap in the genes modulated across the different times. On the other hand, there were several bacterial DEGs that were common across 3 h - 24 h. The maximum number of human DEGs was also observed when bacteria first interacted with cells (3 h) (adjusted p-value < 0.05, and a log_2_(fold change) > 1 or < -1). 140 genes were upregulated, and 175 genes were downregulated at 3 h and in subsequent time points the changes seen were lower, with an increase seen at 12 h (Fig. 2C). For bacterial DEGs, there were 137 upregulated and 149 downregulated genes identified at 3 h, but many of these genes stayed altered until 24 h (Fig. 2C). The top 10 human and bacterial genes (ranked by p adj value), at different times after infection compared to the respective uninfected controls are shown as a heatmap (Fig. 3A, B) and the complete lists of DEGs compared to uninfected controls are included in Table S2. Volcano plots show the distribution of all DEGs at each time point for human and bacterial genes in the control vs infection analysis (Fig. 3C and Fig. S3).

**Figure 2.**
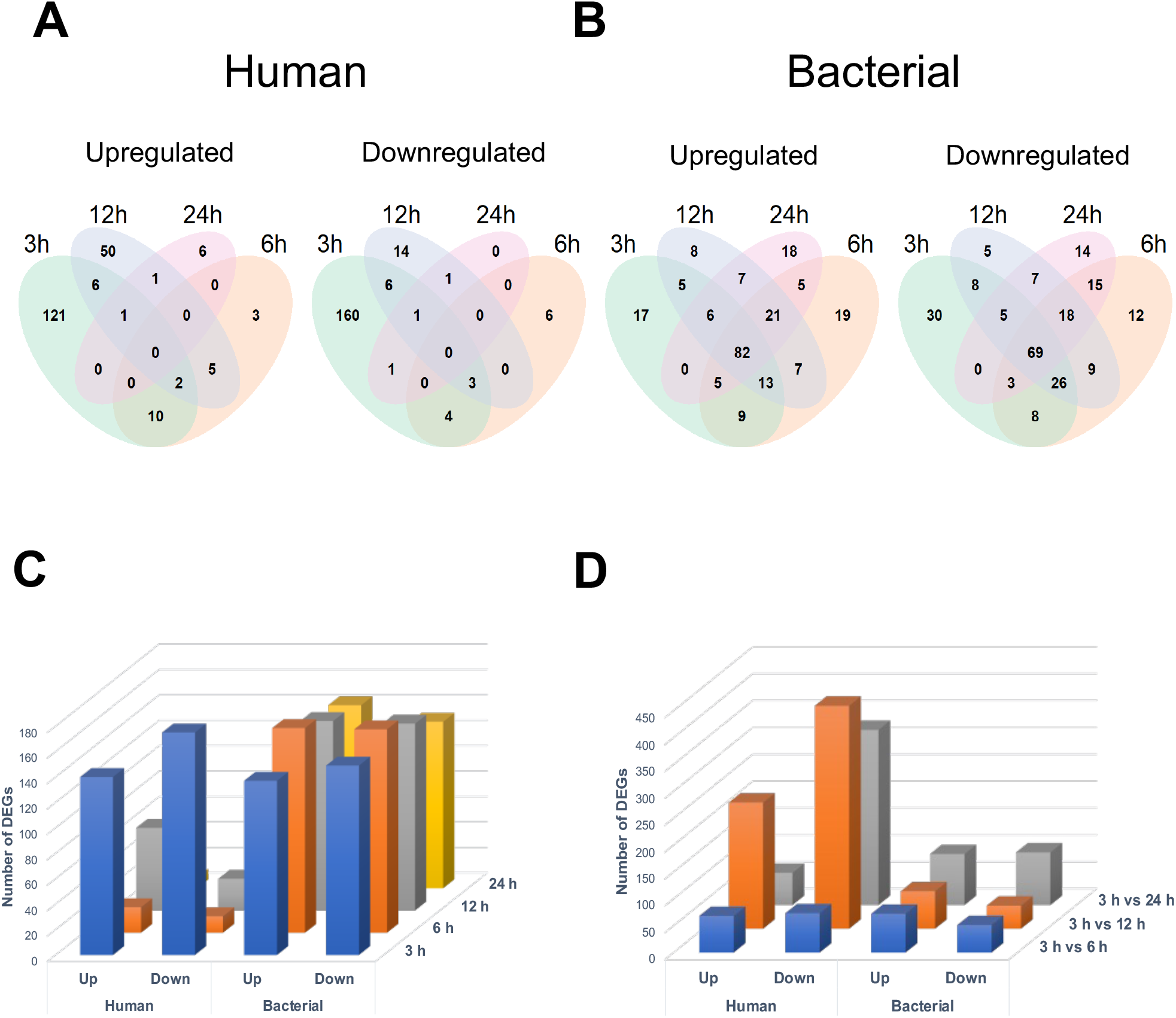
Overall profiles of differentially expressed genes in bacterial and human cells. Venn diagrams of the differentially expressed host (A) and bacterial (B) genes to visualise genes common up- or downregulated between each timepoint. C) Differentially expressed genes (DEGs) at 3, 6, 12 and 24 h post infection, as compared to the relevant uninfected controls. D) Differentially expressed genes between the 3 h infected samples and the other infected samples at 6, 12 and 24 h post infection.

**Figure 3.**
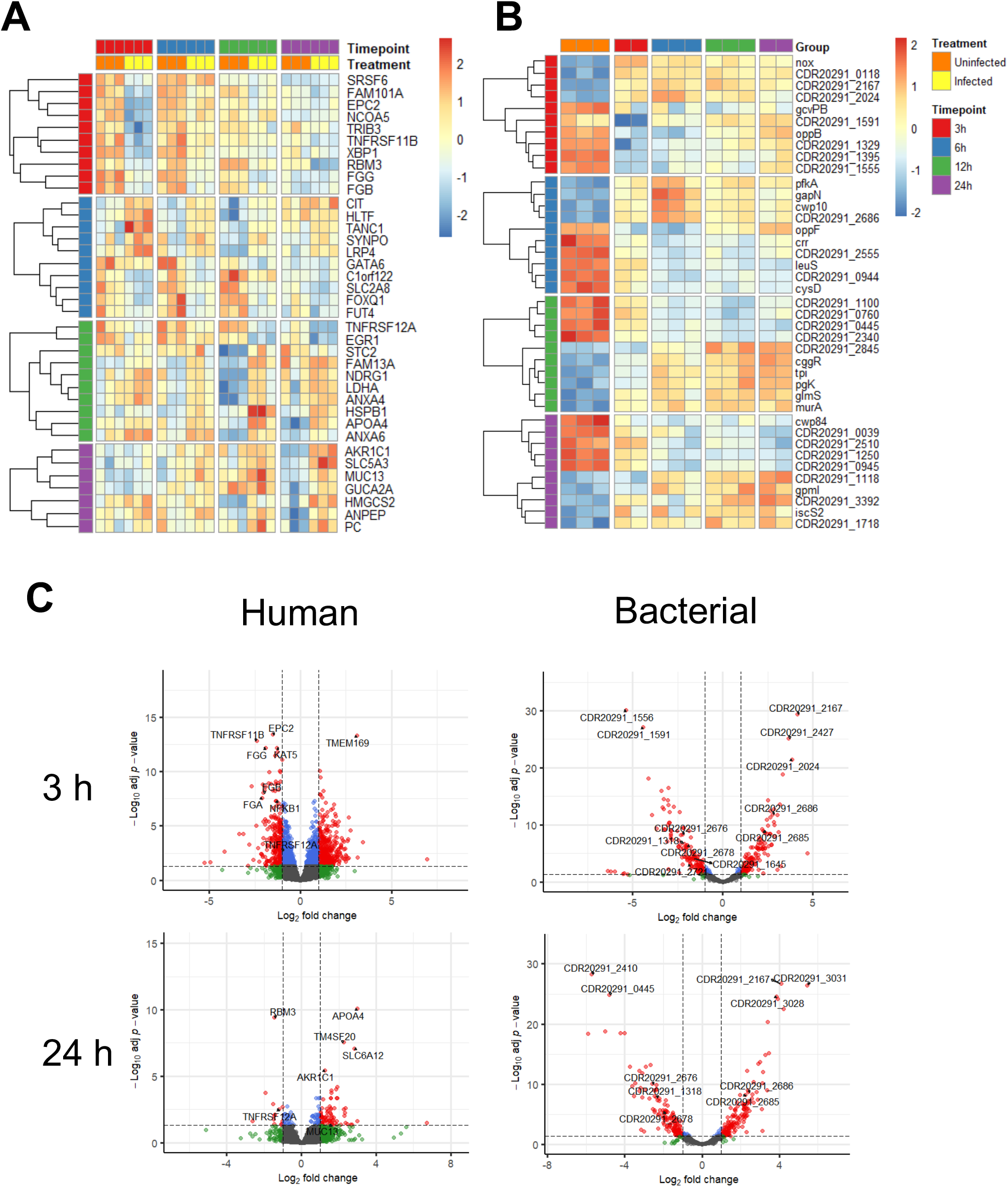
Differentially expressed host and bacterial genes. Heatmaps showing the top 10 most significant differentially expressed genes at each time point as compared to the respective uninfected control for human (A) and bacterial (B) genes C). Volcano plots to visualise the distribution of human and bacterial genes of uninfected controls vs infected samples at 3 h and 24 h after infection. Significant DEGs can be visualised as having a log_2_(fold change) greater than 1 or less than -1 and adjusted p-value less than 0.05 (red dots). Labels and arrows point out selected highly significant DEGs.

A comparison was also made across time points to understand the dynamic changes in gene expression. Time dependent analysis was performed by comparing each time point to 3 h after removal of genes changing in the uninfected controls (Fig. 2D, Table S3); this showed changes occurring over time, with maximal changes (with more genes downregulated) observed between 3 h and 24 h. The top 10 DEGs when compared across different time points 3 h vs 6 h, 3 h vs 12 h, and 3 h vs 24 h, after removing genes changing in the uninfected controls, are included in Table 1. Distribution of DEGs by time-dependent analysis is shown in volcano plots in Fig. S4.

**Table 1.** Top 10 most significantly different human genes changing at different times compared to 3 h after infection

| Time after infection (h) | Gene | log<sub>2</sub>(fold change) | Padj |
| --- | --- | --- | --- |
| 3h vs 6h | BCL6 | -1.4263 | 2.48E-13 |
|  | PPRC1 | 1.461462 | 7.74E-12 |
|  | LCA5 | -1.70836 | 3.39E-09 |
|  | CITED2 | -1.65225 | 2.90E-08 |
|  | JARID2 | -1.00072 | 7.70E-08 |
|  | BEND3 | 1.47559 | 1.28E-07 |
|  | HGNC:9982 | 1.153837 | 1.40E-07 |
|  | EPC2 | 1.230927 | 1.62E-07 |
|  | MYC | 1.188266 | 3.41E-07 |
|  | FZD4 | -1.48643 | 5.72E-07 |
| 3h vs 12h | PHLDB2 | -2.31212 | 8.69E-26 |
|  | NOP16 | 1.871325 | 4.35E-17 |
|  | CDC14A | -2.52359 | 4.35E-17 |
|  | CPLX1 | 2.625903 | 3.00E-16 |
|  | AMOT | -1.63245 | 6.42E-15 |
|  | PDZD7 | -2.25312 | 9.95E-15 |
|  | CITED2 | -2.06608 | 1.69E-14 |
|  | FZD4 | -2.02591 | 2.55E-14 |
|  | LCA5 | -1.89066 | 8.22E-13 |
|  | ZNF608 | -1.73279 | 2.49E-11 |
| 3h vs 24h | GNRH2 | -3.1352 | 7.45E-21 |
|  | ARL4C | -1.8818 | 7.45E-21 |
|  | CCND3 | -2.44675 | 1.67E-18 |
|  | REEP2 | -2.94226 | 2.10E-17 |
|  | MSR1 | -1.98428 | 5.15E-16 |
|  | PDZD7 | -2.24464 | 1.30E-13 |
|  | EHD1 | -1.38567 | 1.09E-11 |
|  | MYLIP | -1.62791 | 1.34E-11 |
|  | BCAS4 | -2.46879 | 1.51E-11 |
|  | VDR | -1.66743 | 4.92E-11 |

### Host responses to *C. difficile* attachment and multiplication

Among the large number of mammalian genes induced in infected cells compared to uninfected controls, one of the most upregulated genes at 12 h and 24 h post infection was apolipoprotein A-IV (*APOA4*), an apolipoprotein primarily produced in the small intestine (Fig. 4A)(32). The gene for mucin 13 (*MUC13*), a transmembrane mucin glycoprotein which is highly expressed on the surface of mucosal epithelial cells in the small and large intestines (33, 34) was upregulated [log_2_(fold change) >1] at 6, 12 and 24 h post infection, but this upregulation was only statistically significant at 24 h post infection. Interestingly, the low-density lipoprotein receptor-related protein 4 (*LRP4*), a protein involved in the inhibition of Wnt signalling (35), was significantly upregulated at 3 h and 12 h after *C. difficile* infection (Fig. 4A). The frizzled-4 protein (*FZD4*), a receptor involved in the Wnt/β-catenin canonical signalling pathway, was significantly upregulated at 3 h after infection, although it did not show any changes in expression at other timepoints (Fig. 4A).

**Figure 4.**
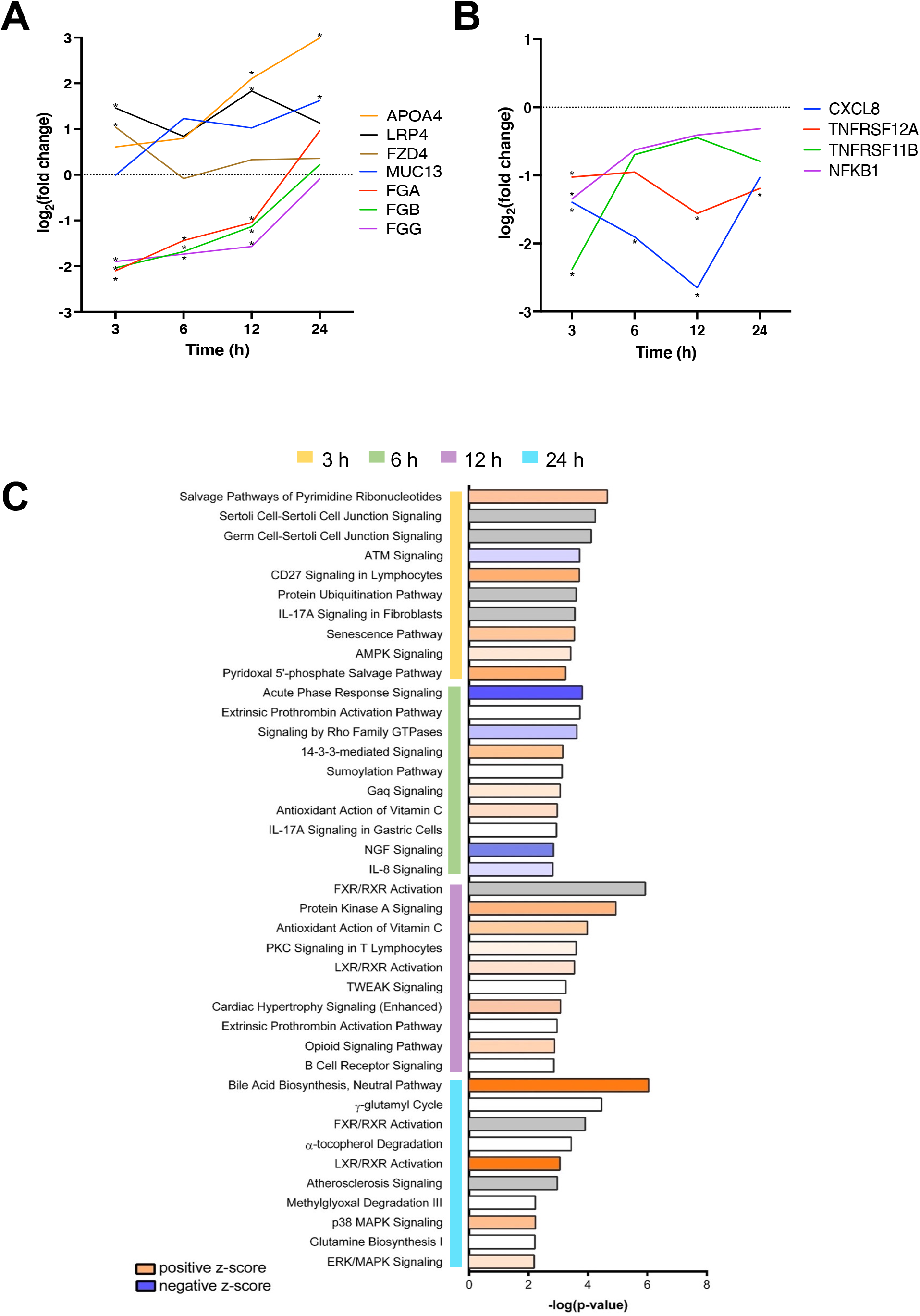
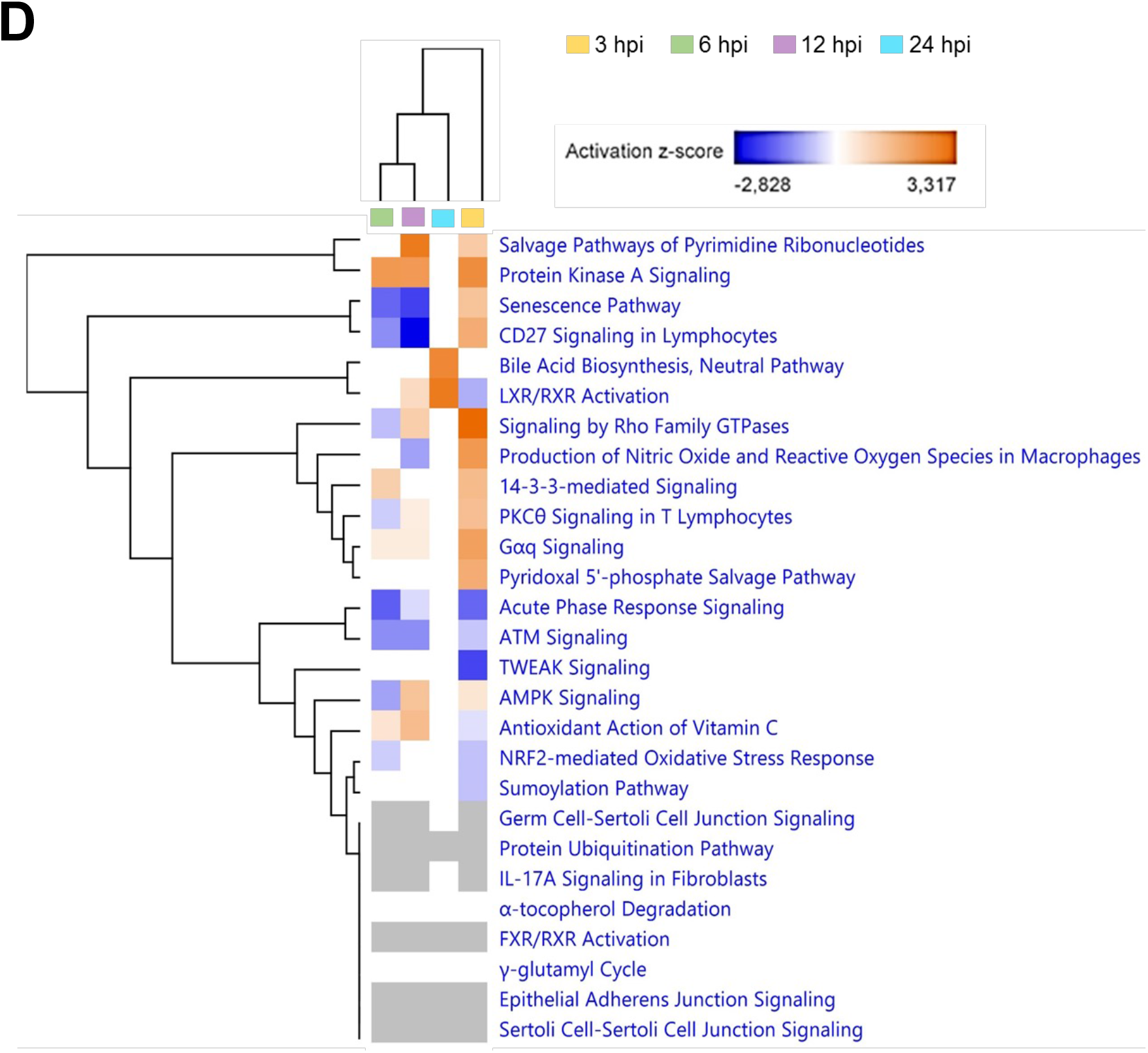
Host pathways and single gene expression profiles during infection. Host single gene expression profiles of selected immune response (**A**) and other interesting genes (**B**) differentially expressed during *in vitro C. difficile* infection. Log_2_(fold change) of selected genes calculated between the expression levels of uninfected controls and infected samples at each timepoint. All selected genes were significantly differentially expressed in at least one timepoint, indicated with an asterisk. (**C**) The z-scores of the top 10 most significantly differentially expressed pathways at each timepoint. (**D)** Heat map of top significantly enriched pathways. The up- (orange) or downregulation (blue) of each pathway is indicated by the z-score. Hierarchical clustering was used to indicate the distances in the gene expression profiles between the timepoints.

One of the most significantly differentially expressed genes at 3 h after infection was tumor necrosis factor receptor superfamily member 11B (*TNFRSF11B*), which encodes osteoprotegerin, a receptor which binds to TNF-related apoptosis-inducing ligand (TRAIL) that was downregulated with a log_2_(fold change) of -2.45 and adjusted p-value of 1.12E-14 (Fig. 4B)(36) Tumour necrosis factor receptor superfamily member 12A (*TNFRSF12A*), known as fibroblast growth factor-inducible immediate-early response protein 14 (Fn14) was significantly downregulated at 3, 12 and 24 h post infection and the nuclear factor kappa B subunit 1 (*NF-κB1*), a gene involved in the inhibition of the NF-*κ*B signalling pathway and the regulation of MAPK signalling was also significantly downregulated after 3 h of infection (Fig. 4B) (37). Surprisingly, the gene for IL-8 (*CXCL8*), a chemokine, strongly associated with *C. difficile* infection was significantly downregulated at 3, 6 and 12 h post infection (31, 38). Also among the highly downregulated genes at 3, 6 and 12 h post infection were the fibrinogen alpha, beta and gamma chain genes (*FGA, FGB* and *FGG*). However, no differential expression was observed at 24 h after infection (Fig. 4B). Thus, several interesting genes that are involved in a range of cellular pathways are regulated during the initial phases of *C. difficile* infection.

Comparing the changes between 3 h and different times after infection (Table 1), there are several genes like *FZD4, LPR4* and *MUC13* which were upregulated compared to the uninfected controls were also upregulated over time. Additionally, other mucin genes like *MUC12* and other FZD proteins (FZD5, FZD7) were upregulated at 12 h and 24 h when compared to 3 h. Matrix metalloprotease-1 (*MMP1*), which encodes for an interstitial collagenase was upregulated when compared between 3 h and different times, although when compared to control uninfected samples, this gene was significantly downregulated at each time point.

To investigate cellular processes and pathways which were modulated during *C. difficile* infection, a pathway enrichment analysis was conducted. Overall, the most significantly modulated pathways included the neutral bile acid synthesis pathway (24 h), the farnesoid X receptor (FXR)/retinoid X receptor (RXR) activation pathway (39), which has a role in the metabolism of bile acids (12 h) and salvage pathways of pyrimidine ribonucleotides, which are responsible for nucleotides recycling from DNA and RNA breakdown (3 h) (Fig. 4C and D). At 3 h, the salvage pathways of pyrimidine ribonucleotides had a positive z-score, suggesting a potential activation during infection, despite some of the genes in the pathway being downregulated. Some other interesting, enriched pathways identified for 3 h post infection, included Sertoli and germ cell-Sertoli cell junction signalling, CD27 signalling in lymphocytes, IL-17A signalling in fibroblasts and 5′-AMP-activated protein kinase (AMPK) signalling. At 6 h after infection the acute phase response signalling, which is involved in the alterations of gene expression and metabolism in response to inflammatory cytokines, had a negative z-score, indicating that this enriched pathway is potentially inhibited during *C. difficile* infection. Other pathways enriched in the data from 6 h post infection, include the extrinsic prothrombin activation, signalling by Rho GTPases, IL-17A signalling in gastric cells, and IL-8 signalling pathways. At 12 h post infection the most enriched canonical pathways were the protein kinase C (PKC) signalling in T lymphocytes, TNF-like weak inducer of apoptosis (TWEAK), extrinsic prothrombin activation pathway and B cell receptor signalling, and at 24 h, enriched pathways include FXR/RXR and LXR/RXR activation, p38 MAPK and extracellular signal-regulated kinase (ERK)/MAPK signalling pathway. Comparing the different time points, we could identify several pathways that are predicted to be activated in all time points post infection, including the protein kinase A, IL-17 and cell junction signalling pathways (Fig 4D).

### Bacterial responses to *C. difficile* infection

There were several interesting differentially expressed *C. difficile* genes identified in this study which have roles across many cellular processes, including virulence, colonisation, and metabolism. Several genes were upregulated at all timepoints, when compared to the *in vitro* culture control, but they were mostly stress associated genes. There were some cell wall associated proteins like *cwp10* (CDR20291_2685), CDR20291_0184 (putative cell wall hydrolase), and CDR20291_2686 which were highly upregulated at all timepoints after infection (Fig 5A). Interestingly, the *cwp84* (CDR20291_2676), a cell wall hydrolase required for the formation of the S-layer was significantly downregulated at all timepoints. *slpA* (CDR20291_2682), the precursor for the S-layer proteins, was also significantly downregulated at 6 h post infection (Fig. 5A). Several other cell-well proteins, such as *cwp66* (CDR20291_2678), a cell wall protein suggested to play a role in adhesion to Vero cells (14), *cwp13* (CDR20291_1645), *cwp14* (CDR20291_2624), *cwp17* (CDR20291_0892), *cwp18* (CDR20291_0903), *cwp19* (CDR20291_2655), and *cwp20* (CDR20291_1318) were downregulated at all timepoints after infection. *codY*, a global transcriptional regulator (40) (CDR20291_1115) was significantly downregulated at 6 h, 12 h and 24 h after infection and *fliC* (CDR20291_0240) at 12 h post infection. Proline-Proline endopeptidase 1 (PPEP-1/CDR20291_2721), an important secreted zinc metalloprotease which cleaves cell-surface associated collagen binding proteins and host extracellular matrix proteins was also significantly downregulated at 3 h, 12 h and 24 h post infection, and is investigated further in this study (11, 41) (Fig. 5B).

**Figure 5.**
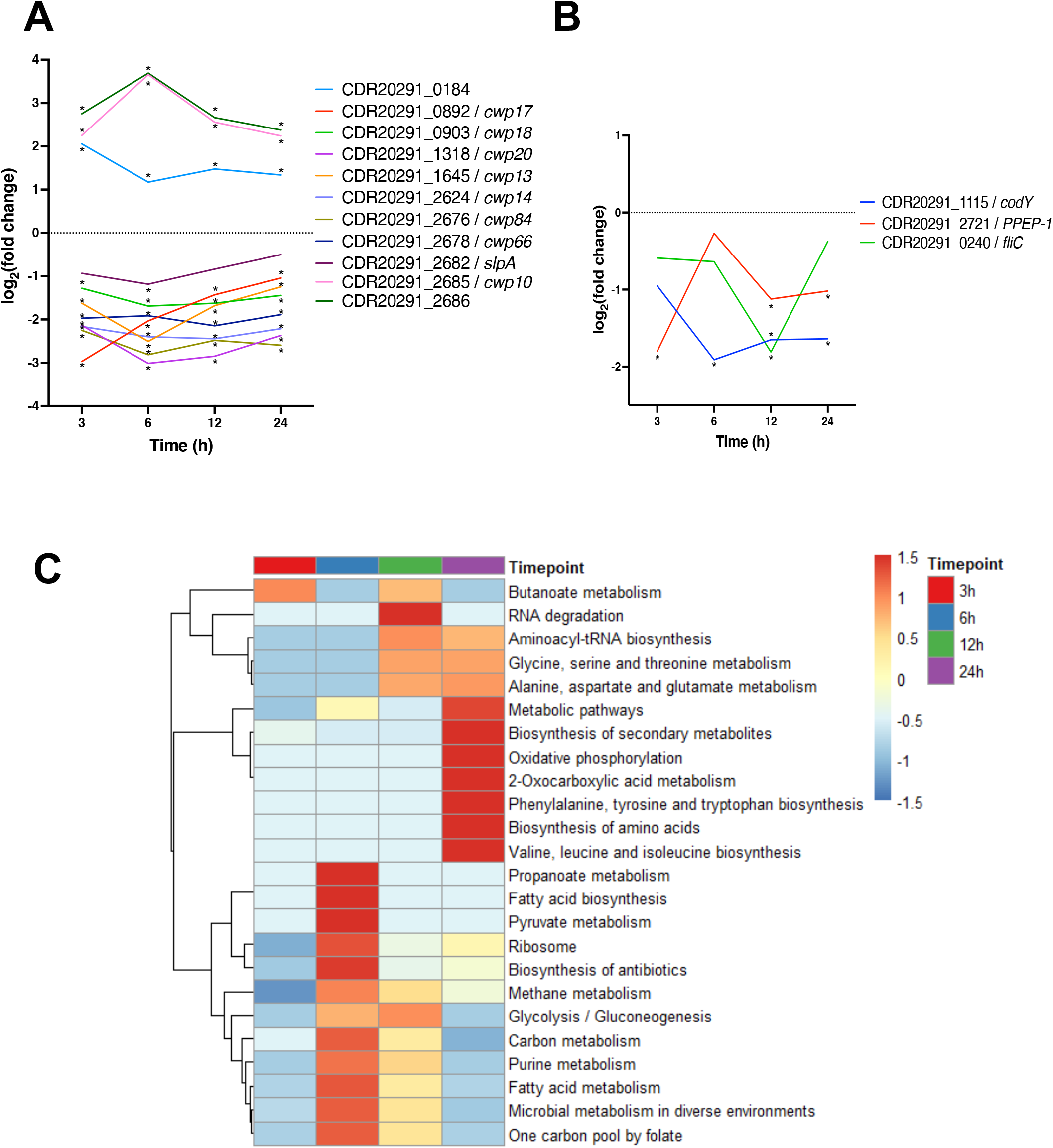
Bacterial genes and pathways changing during infection. Single gene expression profiles of selected C. difficile virulence associated genes (**A)** and *C. difficile* cell-surface proteins (**B**) at different timepoints of infection. Statistical significance indicated with an asterisk. **(C)** Heatmaps of the KEGG pathway functional enrichment analysis of significantly differentially expressed genes during infection. Colours are representative of the enrichment significance (-log_10_(FDR)) and the dendrograms illustrate the hierarchical clustering of samples and pathways.

Bacterial genes changing between 3 h and other times after infection were compared and there were many metabolic pathway genes, and few cell wall proteins changing over time (Table 2). Like the host genes, we see some bacterial genes that are downregulated in relation to the uninfected control, but upregulated when compared across time (e.g., CDR20291_0892).

**Table 2.** Top 10 most significantly different bacterial genes changing at different times compared to 3 h after infection

| Time after infection (h) | Gene | log<sub>2</sub>(fold change) | Padj |
| --- | --- | --- | --- |
| 3h vs 6h | serS1 | 3.58965355 | 4.00E-14 |
|  | dnaH | -3.6691402 | 7.72E-11 |
|  | CDR20291_2824 | 2.05554368 | 9.22E-10 |
|  | recD | 2.51963688 | 2.18E-09 |
|  | pyc | 2.24137439 | 4.87E-09 |
|  | CDR20291_0755 | 2.15269859 | 5.59E-09 |
|  | clpC | -2.1494402 | 5.59E-09 |
|  | CDR20291_0372 | 3.56674282 | 7.00E-09 |
|  | CDR20291_1118 | 2.67543257 | 7.00E-09 |
|  | gltX | 2.68921553 | 1.35E-08 |
| 3h vs 12h | CDR20291_1591 | 4.25801398 | 9.87E-24 |
|  | CDR20291_2845 | 3.6460206 | 3.11E-20 |
|  | CDR20291_2824 | 2.15865642 | 2.49E-12 |
|  | rbo | 2.38713598 | 9.17E-12 |
|  | rbr | 2.26749276 | 2.05E-11 |
|  | CDR20291_1118 | 2.80879361 | 6.36E-11 |
|  | CDR20291_0756 | 2.49678131 | 1.27E-10 |
|  | CDR20291_0760 | -3.1958283 | 1.65E-09 |
|  | CDR20291_2487 | 3.19721177 | 1.26E-08 |
|  | CDR20291_0318 | 2.59373112 | 1.73E-08 |
| 3h vs 24h | CDR20291_1591 | 4.4613226 | 1.81E-26 |
|  | CDR20291_2487 | 4.97667554 | 7.12E-22 |
|  | CDR20291_1118 | 3.79570722 | 3.61E-21 |
|  | rbo | 3.04052554 | 3.91E-20 |
|  | etfA1 | 4.77756376 | 4.49E-19 |
|  | CDR20291_0756 | 3.24899191 | 5.34E-19 |
|  | hadA | 3.35378085 | 8.00E-19 |
|  | hadC | 4.3369607 | 2.01E-18 |
|  | CDR20291_2845 | 3.39039969 | 6.75E-18 |
|  | etfB1 | 4.28467697 | 9.39E-18 |

To investigate bacterial pathways modulated during *C. difficile* infection, we performed a functional enrichment analysis of Kyoto Encyclopaedia of Genes and Genomes (KEGG) pathways. The up- or downregulated genes at each timepoint were entered into the STRING database online tool to identify functional pathways enriched during *C. difficile* infection. There were many interesting significantly modulated pathways identified in this analysis and most of these pathways were associated with the biosynthesis or metabolism of various substrates (Fig. 5C). The metabolic pathways of purines, pyrimidines, carbon, and methane were significantly upregulated during *C. difficile* infection. The biosynthesis of antibiotics, secondary metabolites and gluconeogenesis pathways were also upregulated during *C. difficile* infection (Fig 5C). The metabolic pathways of nitrogen, 2-oxocarboxylic acid, cysteine, methionine, butanoate, glycine, serine, threonine, propanoate, phenylalanine, fatty acid, pyruvate, carbon, glyoxylate and dicarboxylate and the biosynthesis pathways of amino acids, secondary metabolites, aminoacyl-tRNA and antibiotics were significantly downregulated during *C. difficile* infection (Fig. 5C). Some pathways, such as carbon metabolism and biosynthesis of secondary metabolites were identified as both up- and downregulated, suggesting that these pathways were modulated, where some genes were upregulated, and others were downregulated.

### Pyrimidine ribonucleotides were modulated during infection

The most significantly modulated host pathway at 3 h post infection was the salvage pathways of pyrimidine ribonucleotides which had a positive z-score, indicating that it was upregulated during infection. The normalised gene expression values of genes involved in the salvage pathways of pyrimidine ribonucleotides were plotted on a heatmap and hierarchical clustering was applied to group genes and samples with similar gene expression profiles (Fig 6A). In general, most of the samples from the four infected groups were clustered together more closely than the uninfected groups, although the 24 h uninfected controls were clustered more closely to the infected sample groups than the other uninfected controls. At 3 h post infection, there was an upregulation of 12 genes and downregulation of 13 genes which are shown in Fig. S5 in the context of this pathway. At the other three timepoints there were also modulation of the genes involved in the salvage pathways for pyrimidine ribonucleotides, including cytidine deaminase which was significantly downregulated at all timepoints (Fig. 6A).

**Figure 6.**
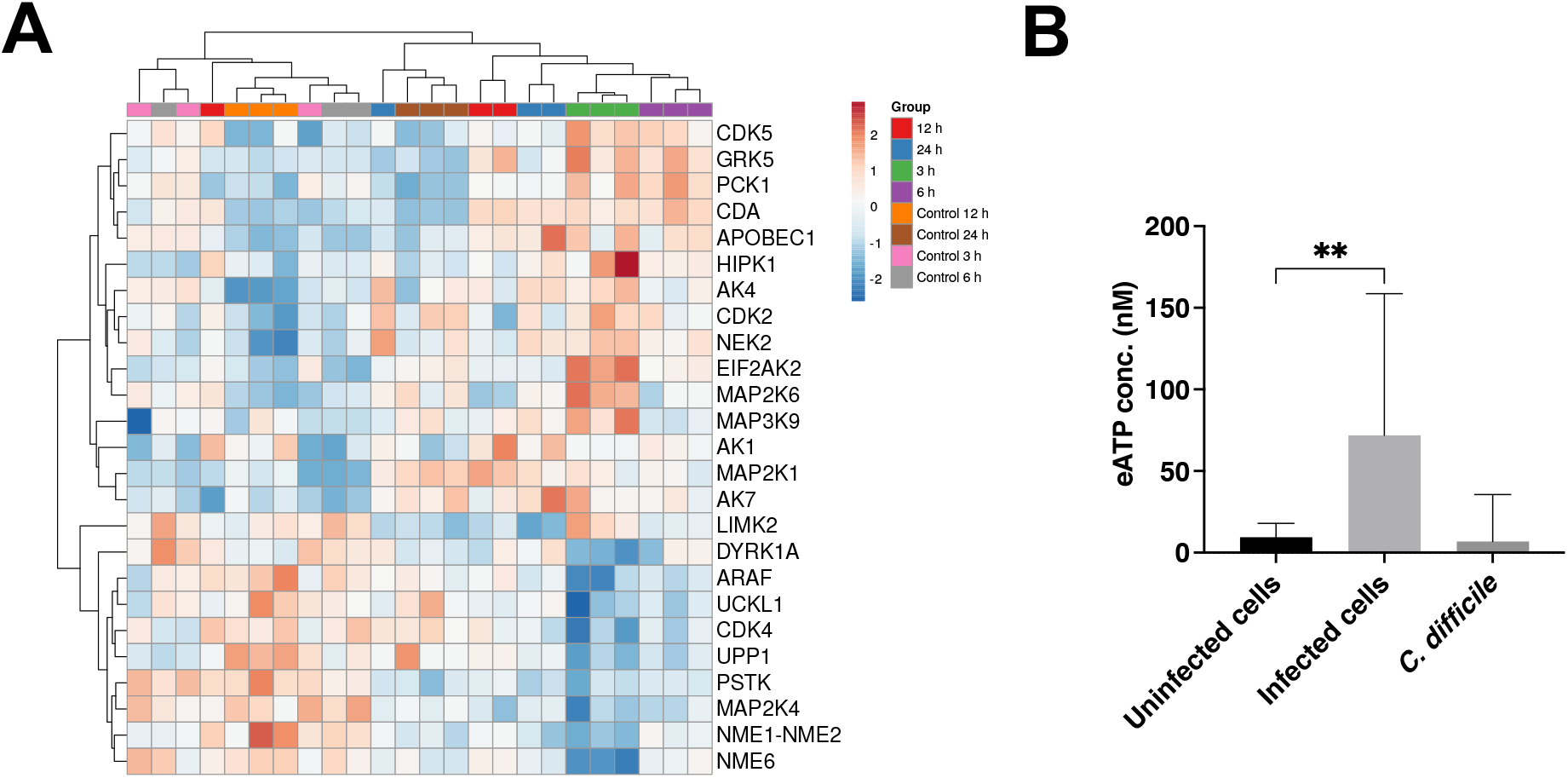
The differential expression of genes involved in the salvage pathways of pyrimidine ribonucleotides. **A)** Heatmap of the genes involved in the salvage pathways of pyrimidine ribonucleotides at 3 h post infection. Regularised logarithmic transformation was applied to the normalised counts from DESeq2. The z-score was calculated to scale the data. The Euclidian distance was used to calculate the gene expression distances between samples, and average linkage was used to cluster the rows and columns. **B)** ATP release assay from uninfected cells, *C. difficile*-infected cells or a bacterial inoculum incubated in pre-reduced DMEM-10 for 4 h under anaerobic conditions. Bars are representative of mean values from 9 technical replicates from 2 biological replicates with standard deviation error bars. **p = 0.007, quantified by paired t-test.

In addition to the salvage pathways of pyrimidine ribonucleotides, there were several other significantly enriched pathways at 3 h after infection with a role in the biosynthesis or metabolism of nucleotides including the nucleotide excision repair pathway, pyrimidine deoxyribonucleotides de novo biosynthesis I, purine nucleotides degradation II (aerobic) and adenosine nucleotides degradation II. Also, as noted from the bacterial gene analysis, purine and pyrimidine metabolic pathways were also significantly differentially expressed in *C. difficile*. To investigate if extracellular nucleotides are involved in infection, we measured extracellular ATP (eATP) an interkingdom, purinergic signaling molecule which is an energy source for cells. A luminescence-based ATP release assay, where ATP is required for the conversion of luciferin into oxyluciferin, was used to quantify the amount of eATP released during *C. difficile* infection *in vitro*; the quantity of luminescence was directly proportional to the concentration of ATP in the reaction. ATP concentrations in supernatants from uninfected and *C. difficile*-infected intestinal epithelial cells were compared. At 4 h post infection, there was significantly more ATP released into the supernatant from *C. difficile*-infected intestinal epithelial cells compared to the uninfected cells (Fig. 6B). Interestingly, there was also a substantial amount of ATP released from the bacterial inoculum (Fig. 6B). This suggests that *C. difficile* may produce some of the eATP in the supernatant from infected cells.

### *∆PPEP-1* may play a role in *C. difficile* colonisation of an intestinal epithelial layer

PPEP-1 is a secreted proline-proline endopeptidase which cleaves collagen-binding adhesins on the *C. difficile* cell surface, and mammalian proteins like fibrinogen (11, 41), although its role during infection is not clear. The gene which encodes PPEP-1 was significantly downregulated at 3, 6 and 24 h after infection, which may result in reduced cleavage of cell-surface associated adhesins and consequently impact formation of biofilm formation and/or bacterial adhesion to host cells. WT and *∆PPEP-1* were first compared for their ability to form biofilms *in vitro* using previously described methods (42). There was no significant difference observed in the quantity of biofilm biomass between WT and *∆PPEP-1* as determined by crystal violet staining (Fig. S6), suggesting that PPEP-1 is not directly involved in *C. difficile* biofilm formation.

To investigate the role of PPEP-1 in *C. difficile* colonisation, intestinal epithelial cells were infected with WT or *∆PPEP-1* in the VDC system, and the number of bacteria attached to the epithelial cells was quantified by CFU counts. A higher number of *∆PPEP-1* bacterial cells adhered to the intestinal epithelial layers in VDCs compared to the WT at 3 h and 6 h post infection, while at 24 h post infection there was no significant difference in adhesion between the WT and ∆PPEP-1 (Fig. 7A). Overall, while a low percentage of WT bacterial cells from the inoculum adhered to the epithelial cell layer at 6 h post infection, there was a significantly higher percentage of *∆PPEP-1* bacterial cells adhered to the epithelial cells compared to the WT (Fig. 7B). Intestinal epithelial layers infected with WT or *∆PPEP-1* in VDCs were also analysed with confocal microscopy to compare levels of bacterial adhesion. Quantitative analysis of the microscopy images confirmed that there were more *∆PPEP-1* bacterial cells attached to the epithelial cell layers compared to the WT at 6 h and 24 h post infection (Fig 7C). Thus, our data indicate that PPEP-1 is downregulated during the initial infection phase, which enables activation of surface proteins that mediate host cell adhesion.

**Figure 7.**
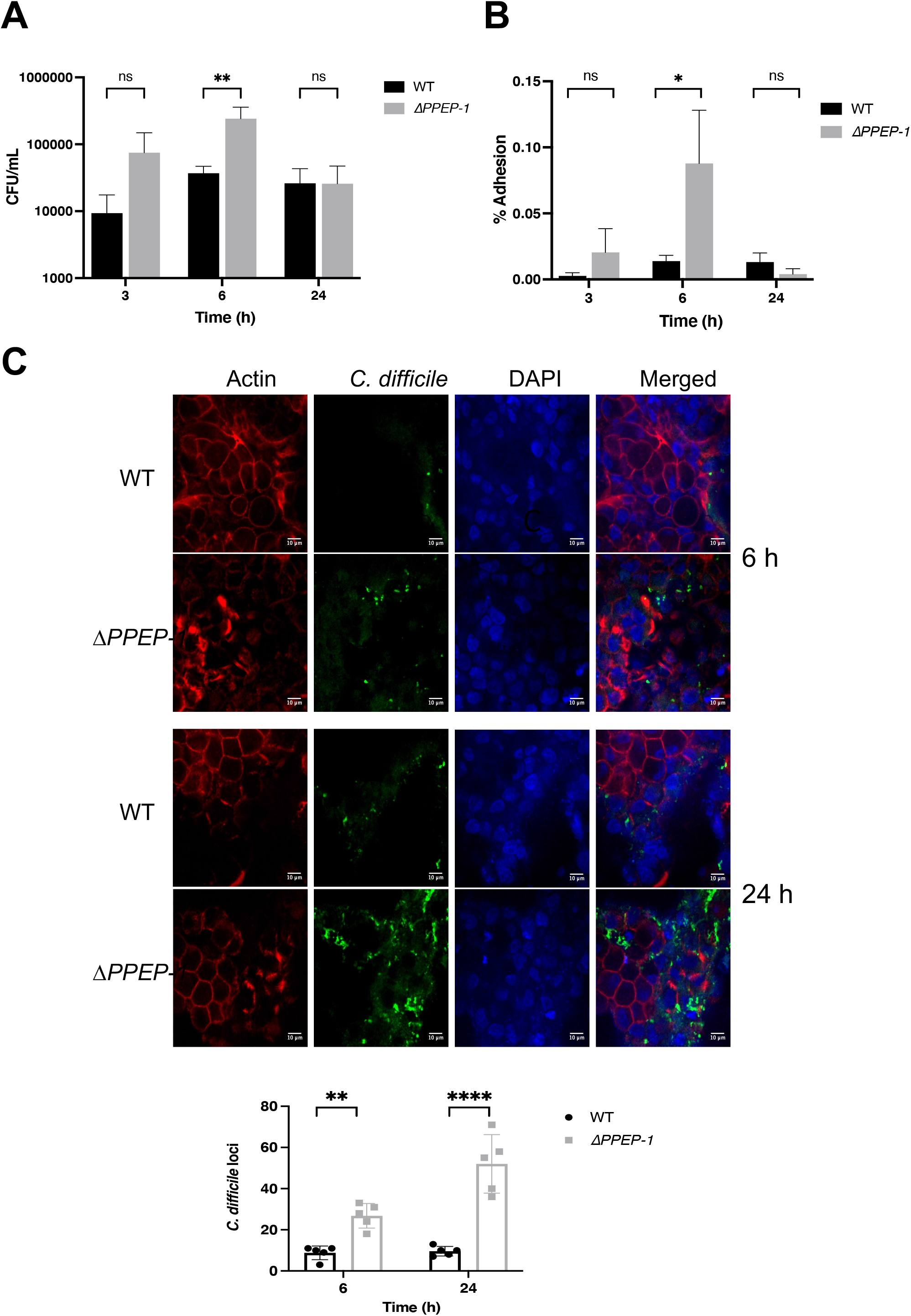
*∆PPEP-1* modulates bacterial adherence to intestinal epithelial cells. A. The number of bacteria attached to intestinal epithelial cells in the VDC system, quantified using CFU counts (n=3) (\*\**p* = 0.003 at 6 h, determined with two-way ANOVA with Sidak multiple comparisons test). B. The percentage of bacterial cells which adhered out of the inoculum, calculated using the equation (CFU of adhered cells/CFU of inoculum) x 100. Statistical significance was tested with an unpaired t-test, \**p* = 0.0342. Bars are representative of the mean +/- standard deviation. C) Cell layers were stained with phalloidin (red), DAPI (blue) and anti-*C. difficile* antibody (green) and imaged with confocal microscopy. *C. difficile* cells adhered to the epithelial cell layers were counted. Statistical significance tested by two-way ANOVA, \*\**p* = 0.0051, \*\*\*\**p* < 0.0001. Data representative of 3 biological replicates with 5 technical replicates/fields analysed. Bars are representative of the mean +/- standard deviation.

## Discussion

*C. difficile* adhesion to the gut mucosa is crucial for the establishment of disease, yet we lack a comprehensive understanding of the host and bacterial factors required for colonisation. Host-pathogen interactions of *C. difficile* infection, particularly in the initial phases of infection, have been largely understudied, mainly due to a paucity of controlled *in vitro* cellular models of infection. Here we report a transcriptomic analysis of epithelial cells infected by *C. difficile* in an *in vitro* gut model which supports co-culturing of anaerobic *C. difficile* with host cells. To our knowledge, this is the first dual RNA-seq study which has been conducted with an anaerobic microbe, in the presence of human gut epithelial cells. Our data show that several host pathways were altered during the initial infection phase, including pyrimidine salvage pathways, modulation of cell wall associated bacterial proteins, and a secreted protease, PPEP-1, which we demonstrate to have a role in mediating adhesion of *C. difficile* to epithelial cells.

The analysis of transcriptomic responses to infection can greatly improve our understanding of the molecular processes which facilitate an infection. Dual RNA-seq is a relatively recent methodology that enables the resolution of the global transcriptomic profiles of both host and pathogen during infection. Unlike probe-based methods, dual RNA-seq sequences both coding and noncoding transcripts which allows the detection of novel and hypothetical genes (43). Transcriptomic responses of numerous bacterial pathogens and infected eukaryotic cells have been resolved with dual RNA-seq, including uropathogenic *Escherichia coli* (UPEC) (44), *Mycobacterium tuberculosis* Bacillus Calmette–Guerin (BCG) (45), *Haemophilus influenzae* (46) and *Salmonella enterica* serovar Typhimurium (47) and have successfully identified new host and bacterial factors which are required to facilitate infections. For example, dual RNA-seq of *Salmonella typhimurium*-infected HeLa cells identified the bacterial small RNA, PinT as a regulator of genes required for intracellular survival and virulence. PinT was also reported to modulate the JAK/STAT signalling pathway in the host cells (47).

Previous transcriptomic studies have focused on *C. difficile* toxin effects on host cells. Inflammation-associated genes (such as C3, CCXL1, CXCL10, DUSP1 and EGR1) played a key role in the response to Tcd1, Tcd2 and a combination of both toxins (27). In a similar study investigating whole-tissue transcriptional responses in *C. difficile* R20291, the IL-1 cytokine family member, IL-33, was significantly upregulated during *C. difficile* infection (19). A recent study by Fletcher *et al*. examined the host transcriptomic responses to WT and *ΔtcdR C. difficile* in a mouse model, although only a targeted set of genes involved in immune responses were assessed using Nanostring analysis (28). Gene Ontology (GO) pathways which were differentially expressed during toxin-induced inflammation included the regulation of the inflammatory response, peptide secretion and proteolysis. In terms of human transcriptomic responses to *C. difficile*, Caco-2 cells were infected with *C. difficile* for 30, 60, and 120 mins in anaerobic conditions and differentially expressed human genes were involved in a range of metabolic processes such as nucleic acid, protein and lipid metabolism, cell organisation and biosynthesis, transport, cell communication, signal transduction, and transcription (38). In this study, we compared the transcriptomes of *C. difficile-*infected epithelial cells to uninfected cells at different times over 24 h post infection, with cells oxygenated from the basolateral side, while exposed to anaerobic conditions on the apical side. It was evident from the changes seen in the control cells over time and the clustering of data, although cells looked morphologically healthy and in an intact layer, cells were exposed to mild stress perhaps due to the apical exposure of anaerobic gas, particularly by 24 h (31). However, distinct profiles were seen at different times after infection, with maximal changes occurring at 3 h, when the cells were first exposed to the bacteria (Fig. 3). While several metabolic pathways were seen to be modulated upon infection, as reported previously (38), it was interesting to note an induction of the LRP and the Frizzled proteins, which are colonic receptors for *C. difficile* TcdA and TcdB (35, 48). During infection within this *in vitro* model, we see very low amounts of toxin A and toxin B produced during the initial 24 h of infection, although this may be sufficient to trigger increased expression (31). Alternatively, the presence of *C. difficile* or other bacterial components may trigger expression of these genes, which leads to the increased cellular binding and uptake of toxins.

Mucins have previously been implicated in *C. difficile* infection. Patients with CDI secrete mucus of altered composition in their feces; with higher levels of the membrane bound mucin, MUC1, and a decreased expression of secretory mucin MUC2, when compared to healthy patient stool (49). It was interesting that the membrane-bound immunomodulatory mucins 12 and 13 are upregulated during infection. MUC13 has been implicated in other microbial infections recently (50), and it would be interesting to explore the role of these immunomodulatory mucins during CDI further. In terms of immune pathways, IL-17 signalling pathways were significantly induced in epithelial cells early in infection, suggesting that this cytokine plays a key role in infection as reported recently (25). There were some immune pathway-associated genes including genes from the TNF receptor superfamily (TNFRSF) which were downregulated during infection, which may suggest that the presence of the pathogen dampens the TNFR-mediated cell signalling. Although transient upregulation in multiple matrix metalloproteinases (MMPs) transcripts was reported previously during toxin-induced intestinal inflammation in mice (28), we see only MMP1 modulated over time in early infection. Overall, it is hard to directly compare the changes induced in the host transcriptome between studies due to differences in the types of host cells and models used, the time post infection and the bacterial strains used.

Pathways involved in salvage of pyrimidine nucleotides were upregulated at all timepoints after infection. The modulation of this pathway during *C. difficile* infection is interesting because extracellular nucleotides have been reported to be regulators of the immune response (51). Bacteria have been reported to produce ATP during growth (52) and the eATP produced by intestinal bacteria in the gut can prevent generation of a protective IgA response against invading bacterial pathogens (53). Our data suggests that *C. difficile* produces a low level of ATP. During infection, ATP produced by bacteria, as well as released by host cells may induce local inflammatory responses in the gut, and nucleotide salvage pathways may be triggered to recycle nucleotides, as well as contain the inflammatory responses.

There have been several studies examining the *C. difficile* transcriptome in different animal infection models. Using a microarray-based approach in a germfree mouse model, Janoir *et al*. reported that majority of the gene expression changes were in metabolism (fermentation and amino acid and lipid metabolism), stress response, pathogenicity, and sporulation, with the toxin genes *tcdA* and *tcdB* moderately upregulated at the later stages of infection (29). A study analysing the nutrients important during *C. difficile* VPI 10463 colonisation using an RNA-seq approach, showed that several metabolic pathways, including carbohydrate, amino acid and fatty acid uptake and metabolism were upregulated early in the process of susceptible host colonisation, with *feoB*, a ferrous iron importer being one of the top upregulated genes at the later stages of infection (54). In a microarray-based analysis of *C. difficile* responses in a pig ligated-loop model infection, upregulation of the *tcdA* was observed 12 h post infection, along with upregulation of colonisation and virulence associated genes, such as CD2830 (*PPEP-1*), CD2592 (fibronectin binding protein), CD2793 (*slpA*), CD1546, CD1208 (hemolysins), and genes involved in the sporulation cascade (30). The present study highlights the modulation of several bacterial metabolic pathways, including fatty acid, amino acid and nucleotide synthesis. The purine and pyrimidine synthesis pathways are initially upregulated, which may indicate an increase in nucleotide production, potentially enhancing an inflammatory response. Additionally, pyrimidine metabolism is known to be essential for *C. difficile* survival (55).

We observed several differentially expressed cell surface proteins, including SlpA, downregulated at all timepoints after infection relative to expression in control medium (DMEM with 10% serum). When compared over time, i.e., 3 h to other timepoints, there are no changes in gene expression observed for *slpA*, but Cwp84 was downregulated. The data suggest that expression of these genes is perhaps dampened when the bacterium is in presence of host cells. With the S-layer being an impervious tightly packed capsule around the bacterium (56), a potential explanation may be that downregulation might enable bacteria to secrete proteases or express surface proteins through S-layer depleted regions of the cell wall. Another gene that was downregulated was *codY*, a global regulator of gene expression and suppressor of several virulence factors, such as toxin production and sporulation (40, 57). This regulator has been reported to be upregulated in other studies although the times and experimental conditions used in these studies were different (38).

A downregulation of the proline peptidase PPEP-1, an enzyme that cleaves bacterial surface adhesins CDR20291_2820 (CD630_28310) and CDR20291_2722 (CD630_32460), was also always seen during infection as reported previously in murine infection (54). *∆PPEP-1* has been previously reported to exhibit an attenuation in virulence in a hamster model (41), although its precise role during infection was unclear. A previous study showed that a PPEP-1 mutant showed enhanced binding to collagen-1 which CD630_28310 was shown to bind to (58). Here, we demonstrate a role for PPEP-1 as a negative modulator of epithelial cell attachment during *C. difficile* infection, suggesting that this enzyme may have a role in the release of *C. difficile* from intestinal epithelial cell layers, through cleavage of cell surface adhesins, aiding *C. difficile* dispersal and penetration of gastrointestinal tissues.

Thus, we describe transcriptional changes occurring in human epithelial cells and in the bacterium during the initial contact between *C. difficile* with host cells within an *in vitro* gut infection model. While the *in vitro* model described in this study mimics the gut environment to some extent, including small amounts of mucus, it lacks the thick complex mucus layer that overlays the gut epithelium. However, this analysis sheds light on the genes and pathways which are modulated during the initial phases of *C. difficile* infection, providing new insight into potential mechanisms of *C. difficile*-host interactions.

## Methods

### Bacterial strains and growth conditions

*C. difficile* 630 and B1/NAP1/027 R20291 (isolated from the Stoke Mandeville outbreak in 2004 and 2005) were primarily used in this study. *ΔPPEP-1* mutant was provided a kind gift from Prof Neil Fairweather, Imperial College London. *Clostridioides difficile* strains were streaked from glycerol stocks and cultured using brain-heart infusion (BHI) agar or broth (Sigma Aldrich, USA) supplemented with 1g/L L-cysteine (Sigma Aldrich, USA) and 5g/L yeast extract (Sigma Aldrich, USA) (BHIS) under anaerobic conditions (80% N_2_, 10% CO_2_, 10% H_2_) in a Don Whitley workstation (Yorkshire, UK).

### Cell culture, media, and conditions

Intestinal epithelial cell line, Caco-2 (P6-P21) from American Type Culture Collection, mucus producing cell line, HT29-MTX (P45-P60), a kind gift from Prof Nathalie Juge, Quadram Institute, Norwich, and intestinal myofibroblast cells, CCD-18co (P10-P20) were used in this investigation. Caco-2 cells were grown in Dulbecco’s modified Eagle medium (DMEM) supplemented with 10% FBS (DMEM-10) (Labtech, UK), and 1% penicillin-streptomycin (10,000 units/mL penicillin, 10 mg/mL streptomycin, Sigma Aldrich, USA). HT29-MTX were grown in DMEM-10 and CCD-18co in Eagle’s Minimum Essential Medium media, both supplemented with 10% FBS, 1% penicillin-streptomycin, 2 mM glutamine, and 1% nonessential amino acids (Sigma Aldrich, USA). All cell lines were maintained in 5% CO_2_ in a humidified incubator at 37°C and free from mycoplasma contamination as determined on a regular basis by the EZ-PCR Mycoplasma kit (Biological Industries, USA). Snapwell inserts (Scientific Laboratory Supplies Ltd, UK) were prepared as described in (31). Briefly, prior to seeding cells, the Snapwell inserts were coated with a 1:1 ratio of rat tail collagen (Sigma Aldrich, USA) and ethanol (VWR Chemicals, USA). Caco-2 and HT29-MTX were mixed in a 9:1 ratio and 2 × 10^5^ cells total was seeded on 12 mm Snapwell inserts (tissue culture treated polyester membrane, Corning, USA) for ∼2 weeks to form a polarised monolayer. Caco-2 and HT29-MTX were seeded on the apical side of the Snapwell insert for 14 days. Prior to infection experiments, the cell culture medium in the Snapwell inserts was replaced with antibiotic-free medium 24 h or 48 h before the start of infection.

### Infection of intestinal epithelial cells in the VDCs

The Snapwell inserts containing the polarised cell layers were placed between the two half chambers of the VDC (Harvard Apparatus, Cambridge, UK) and sealed with the provided clamps. 3 mL DMEM-10 was added to both sides to fill the chamber. As described previously, (31) a single bacterial colony was inoculated in pre-reduced BHIS broth and incubated at 37°C for 16 h in anaerobic conditions. The culture was centrifuged at 5,000 rpm for 5 min (Eppendorf 5810R, Eppendorf, Germany) and bacterial pellet was resuspended in DMEM-10, before being diluted to an OD_600_ of 1.0 and incubated at 37°C in anaerobic conditions for 1 hour. Intestinal epithelial cells were infected with bacterial cultures at an MOI of 100:1 in the apical chamber. This chamber was supplied with anaerobic gas mixture (10% CO2, 10% H2, 80% N2, BOC, UK) and the basolateral compartment with 5% CO2 and 95% air (BOC, UK) at a rate of ∼1 bubble every 5 seconds. At 3 h post infection, the apical media containing the *C. difficile* was removed, the chamber was washed once with PBS and 3 mL fresh pre-reduced DMEM-10 was added. Chambers were incubated for a further 3 – 45 h. The intestinal epithelial cell layers were washed thrice with pre-reduced PBS before being lysed with 1 mL sterile water. The number of cell-associated bacteria were quantified by performing serial dilutions and colony forming unit (CFU) counts from the intestinal epithelial cell lysate, prepared using serial dilutions plated on BHIS agar.

### RNA isolation

Bacterial and human cells were treated with RNAprotect cell or bacterial reagent (Qiagen, Germany) and stored at -80°C for up to 1 week. To extract RNA, samples were thawed and centrifuged at 4°C (Sigma Zentrifugen 1-14K, Germany). Cell pellets were resuspended in 1 mL buffer RLT from the RNeasy mini kit (Qiagen, Germany) with a 1:200 dilution of *β*-mercaptoethanol (Sigma Aldrich, USA). Cells were homogenised using lysing matrix B tubes with 0.1 mm silica beads (MP Biomedicals, USA) in a FastPrep-24 5G (6.5 m/s for 20 seconds with 180 seconds rest on ice for 6 cycles) (MP Biomedicals, USA). RNA was extracted using the RNeasy RNA isolation kit, (Qiagen, Germany), according to the manufacturer’s protocol. A rigorous treatment with TURBO DNase (Thermo Fisher Scientific, USA) was used to remove genomic DNA contamination from RNA samples and clean up was performed using the RNeasy mini kit following the manufacturer’s protocol (Qiagen, Germany). RNA concentrations were quantified using the Qubit RNA BR assay kit (Thermo Fisher Scientific, USA). The RNA quality was examined using the Bioanalyzer RNA 6000 pico kit (Agilent, USA).

### Ribosomal RNA depletion, library preparation and RNA sequencing

An RNase H method for rRNA depletion was used as described by Adiconis *et al*. (2013). The human rRNA single-stranded DNA probes described in this thesis were purchased as oligonucleotides (Integrated DNA technologies, USA). *C. difficile* probes complementary to the 16S, 23S and 5S rRNA sequences were designed as ∼50 bp non-overlapping primers (Integrated DNA Technologies, USA). RNase H rRNA depletion protocol was used as described by SciLifeLab (Sweden). Briefly, oligonucleotide probes were hybridised to RNA samples by incubation with hybridisation buffer (1 M NaCl, 0.5 M Tris-HCl pH 7.4) in a Bio-Rad T100™ thermocycler (95°C for 2 min, temperature decreased -0.1 C/s to 45°C and hold at 45°C). Thermostable RNase H (New England Biolabs, USA) was added, and the reaction was incubated at 45°C for 30 min. Agencourt® AMPure® XP magnetic beads were used to purify RNA. DNase I (New England Biolabs, USA) was used to remove leftover DNA oligonucleotides and Agencourt® AMPure® XP magnetic beads (Beckman Coulter Life Sciences, USA) were used to clean up rRNA depleted samples. Bioanalyzer RNA pico analysis was used to assess the RNA quality and effectiveness of the rRNA depletion (Agilent, USA). The NEBNext® Ultra™ II Directional RNA Library Prep Kit for Illumina® with the NEBNext® Multiplex Oligos for Illumina® (Index Primers Set 1) were used to generate DNA libraries for sequencing (New England Biolabs, USA) as per the manufacturers protocol. Agencourt® AMPure XP magnetic beads (Beckman Coulter Life Sciences, USA) were used for bead-based DNA clean up. Quality of DNA libraries were assessed using a Bioanalyzer High Sensitivity DNA kit (Agilent, USA). Single-end sequencing was performed using a NextSeqR 500/550 High Output Kit v2 (75 cycles) on an Illumina NextSeq™500 system (Illumina, USA). Bacterial controls were sequenced using single-end sequencing with a MiSeq™ v3 cartridge (150 cycles) on an Illumina MiSeq™ system.

### RNA-seq analysis

Raw sequencing files were converted to FASTQ files using the bcl2fastq Linux software package (version 2.20) and the quality of the FASTQ files was assessed using the FastQC Linux package (version 0.11.9). FASTQ files were mapped to the appropriate reference genome (FN545816.1 for *C. difficile* R20291 and GRCh38 for human) using HISAT2 (version 2.1.0). SAM files were converted to sorted BAM files and sorted using the Samtools (version 1.3.1) ‘view’ and ‘sort’ functions. Abundance of each genomic feature was analysed using HTseq-count (version 0.11.2). The data were filtered to only include genes which had > 10 reads in > 10 samples. The DESeq2 (version 1.36.0) R package was used to calculate the differential gene expression profile using a negative binomial distribution model. Differences between groups were considered significantly different where the adjusted *p*-value was below 0.05 and log2(fold change) was greater than 1 or less than -1 for human and bacterial samples. The ‘ggplot2’ R package (versions for R and ggplot2 3.3.6) was used to generate data visualisations, including principal component analysis (PCA) plots, Venn diagrams, etc. Heatmaps were generated using the ‘pheatmap’ R package (version 1.0.12) and volcano plots were generated with the ‘EnhancedVolcano’ R package (version 1.14.0). For visualisation, gene expression data was scaled and centred using the z-score. In hierarchical clustering, row and column clustering distance was calculated with the Euclidean distance and clustered with the average linkage method. All sequencing reads were deposited to the European Bioinformatics Institute ArrayExpress (Accession number E-MTAB-12660).

### Pathway and functional analyses

Pathway and functional analyses were performed using Ingenuity Pathway Analysis (IPA) software (QIAGEN, USA). The lists of differentially expressed genes from the host and bacteria, were uploaded to IPA software and analysed for enrichment of gene annotation terms to obtain a list of canonical pathways potentially modulated during host-bacteria interaction. For these analyses, Fisher’s exact test was used to measure the enrichment significance based on the number of molecules/genes overlapping each identified pathway. Also, sequencing reads were aligned to the *C. difficile* strain 630 genome, analysed for differential expression and analysed for enrichment using the STRING online database (https://string-db.org) to investigate bacterial KEGG pathways, protein-protein interaction networks.

### Immunofluorescent staining and confocal microscopy analysis

Epithelial cell layers infected with *C. difficile* as described above were washed thrice with PBS to remove unadhered bacteria and fixed with 4% paraformaldehyde (PFA) (Alfa Aesar, USA) for 15 min at RT. Cells were permeabilised with 1% saponin (Sigma Aldrich, USA) in 0.3% triton X-100 (Sigma Aldrich, USA) in PBS (Thermo Fisher Scientific, USA) and then blocked with 3% BSA (Sigma Aldrich, USA) in PBS. Rabbit anti-*C. difficile* sera was used as a primary antibody to stain *C. difficile* on infected intestinal epithelial cells (1:500 dilution in 1% BSA in PBS for 1 h at room temperature), followed by goat antirabbit IgG AlexaFluor 488 conjugate antibody (1:200 dilution in 1% BSA in PBS for 1 h at room temperature in the dark) (Cell Signalling Technology, USA) as the secondary antibody. AlexaFluor 647 phalloidin (Cell Signalling Technology, USA) was used at a 1:100 dilution in PBS to stain the actin cytoskeleton. ProLong Gold Antifade Reagent with DAPI (Cell Signalling Technology, USA) was used to stain cell nuclei and seal coverslips. Slides were imaged using a confocal spinning-disk microscope (VOX UltraView, PerkinElmer, USA) with a 40X oil objective and two Hamamatsu ORCA-R2 cameras, by Volocity 6.0 (PerkinElmer, USA). Image analysis was performed using the Fiji software package (version 2.0.0).

## Acknowledgements

We would like to acknowledge the Wellcome Warwick Quantitative Biomedicine Programme Seed grant and the Medical and Life Sciences Research Fund for funding this study and the Midlands Integrative Biosciences Training partnership PhD studentship (1782606) to Lucy Frost. We would like to thank Dr Chrystala Constantinidou at the Warwick Bioinformatics Research Technology Platform for discussions and support with bioinformatic analysis.

## Supplementary figures

**Figure S1** PCA plot of host RNA-seq data to show sample-sample distances of infected samples and uninfected controls at 3, 6, 12 and 24 h. B. PCA plot of bacterial RNA-seq data to show sample-sample distances of infected samples at 3, 6, 12 and 24 h compared to an uninfected control culture. The red circles indicate samples which were removed from the analysis due to high variation from the other replicates. C. PCA plot of bacterial RNA-seq data after the removal of two highly distant samples.

**Figure S2** Heatmaps of host and bacterial transcriptomic data. A. Heatmap of host transcriptomic data to demonstrate the variations in gene expression between the infected samples and uninfected controls at each timepoint. B. Heatmap of bacterial transcriptomic data to demonstrate the variations in gene expression between the infected samples and uninfected controls at each timepoint. Differential gene expression values calculated using the z-score, which is represented with a colour scale, where red indicates a positive z-score, and blue indicates a negative z-score.

**Figure S3** Volcano plots to illustrate the distribution of human and bacterial genes of uninfected controls vs infected samples at 6 h and 12 h after infection. Significantly differentially expressed genes can be visualised as having a log2(fold change) greater than 1 or less than -1 (blue lines) and adjusted p-value greater than 0.05 (red line). Arrows point out selected highly significantly differentially expressed genes.

**Figure S4** Volcano plots to illustrate the distribution of human and bacterial genes of infected samples from 3 h compared with 6, 12 or 24h after infection. Significantly differentially expressed genes can be visualised as having a log2(fold change) greater than 1 or less than -1 (blue lines) and adjusted p-value greater than 0.05 (red line). Arrows point out selected highly significantly differentially expressed genes

**Figure S5** PPEP-1 does not impact biofilm formation

Biofilms were developed over 24 h in BHIS+G media as quantitated by crystal violet staining. Significance tested with one-way ANOVA with multiple comparisons to the WT as a control, p = 0.013. Bars are representative of the mean +/- standard deviation

**Figure S6** A pathway map of the pyrimidine ribonucleotide de novo biosynthesis KEGG pathway. Upregulated genes are coloured in red and downregulated genes are coloured in green.

**Table S1** Total number of reads, percentage of reads aligned to a concatenated dual reference genome or *C. difficile* genome of all sequenced samples.

**Table S2** Excel file with all significant DEGs from all time points compared to control for bacterial and human samples.

**Table S3** Excel file with all significant DEGs from all time points compared to 3 h for bacterial and human (DEGs changing in control samples were removed) samples

