## Supplementary figures and images for "Dual RNA-seq identifies proteins and pathways modulated during *Clostridioides difficile* colonisation"

### Supplemental Figures

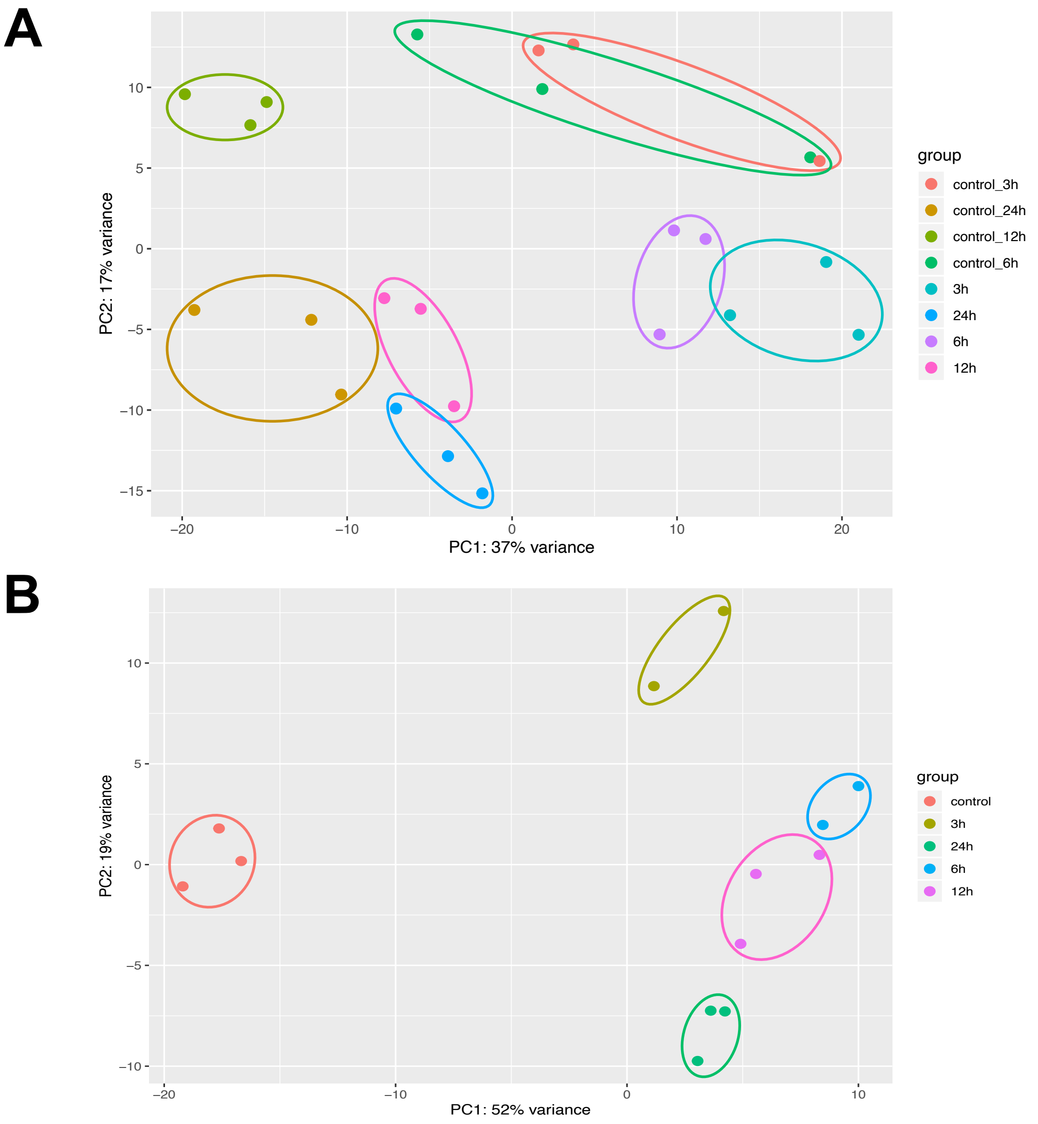

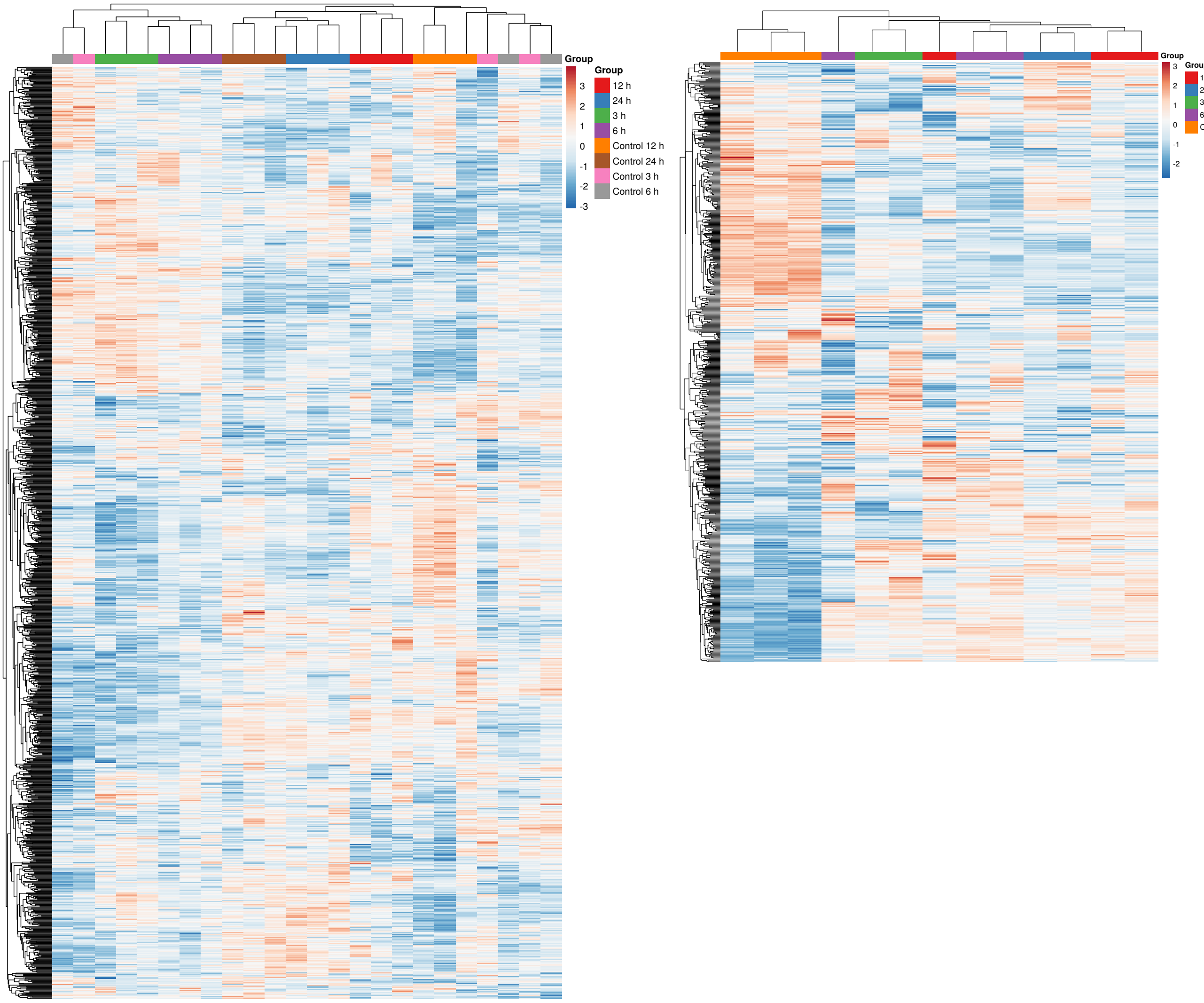

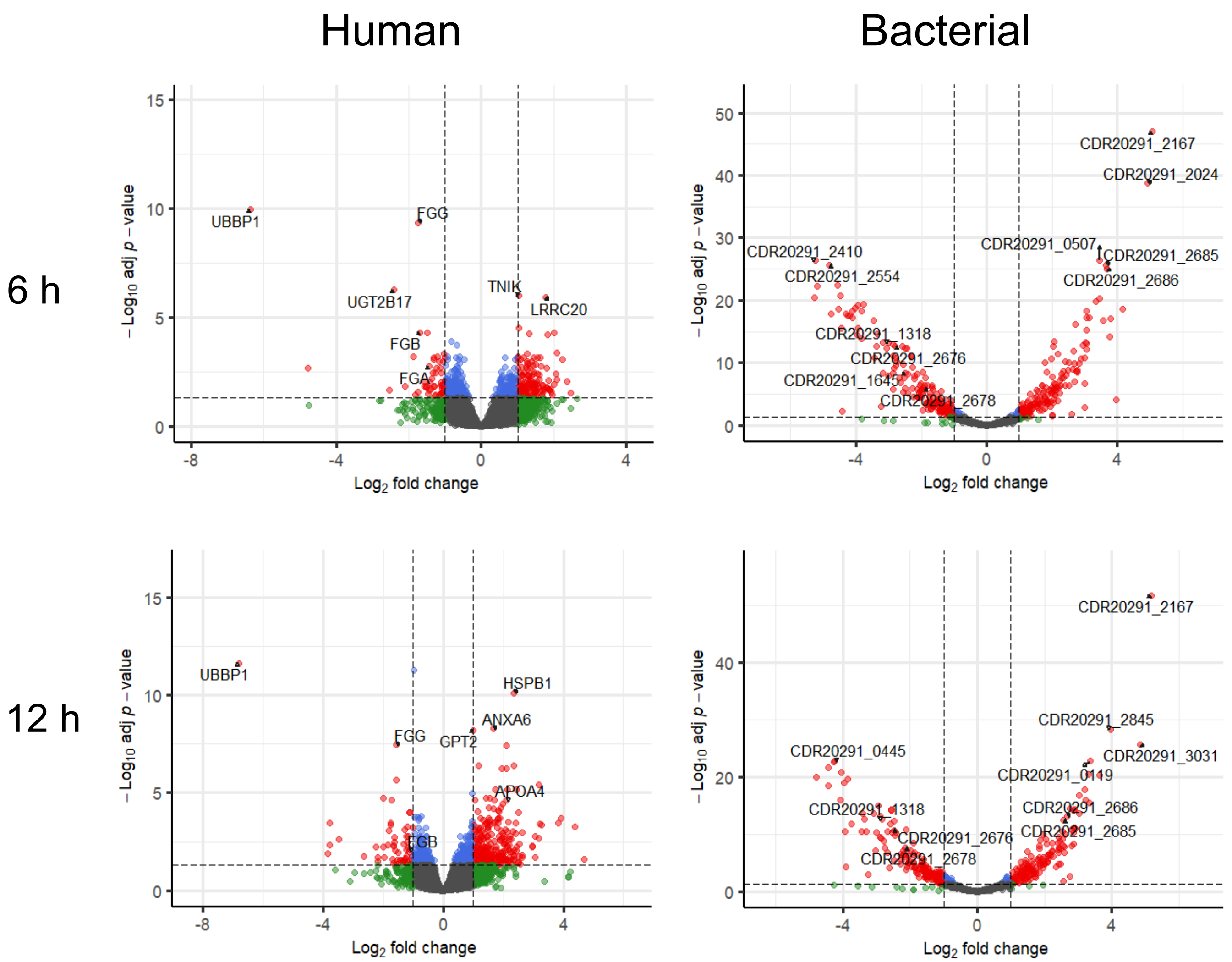

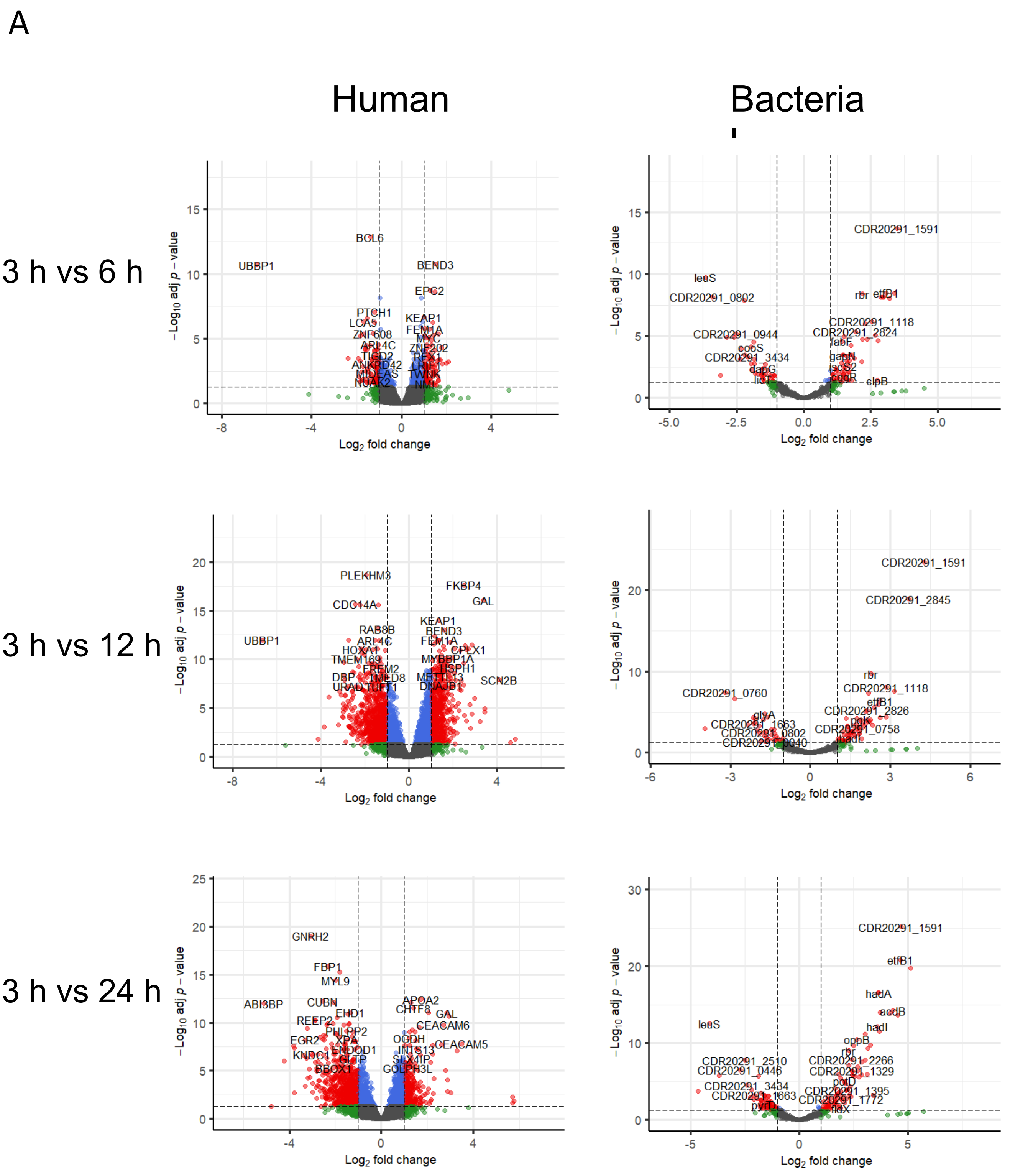

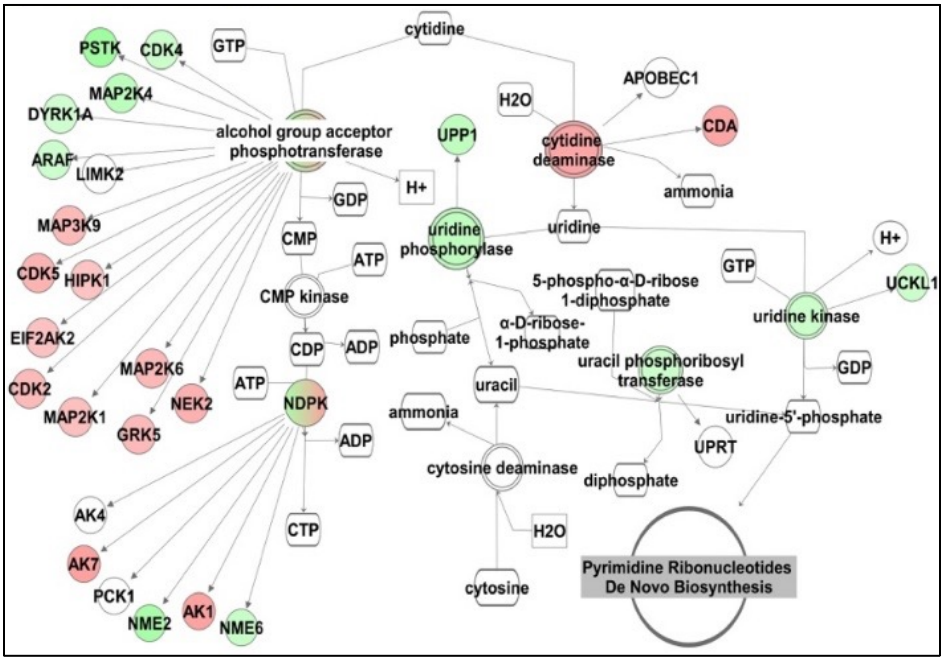

#### Figure S6

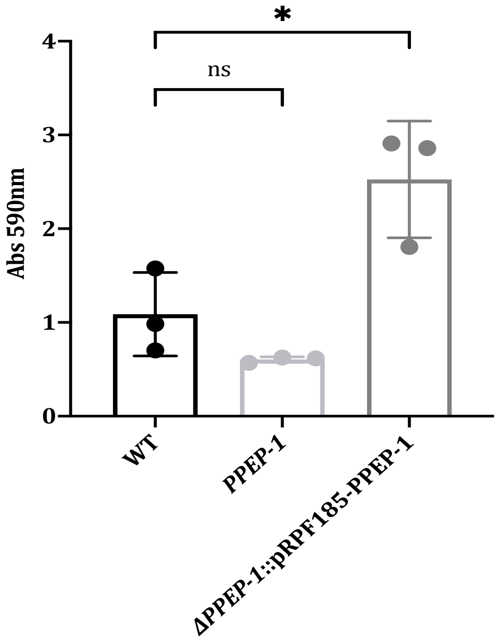
