## Supplementary material for "Dual RNA-seq identifies proteins and pathways modulated during *Clostridioides difficile* colonisation": Table S1

**Table S1** **Total number of reads, percentage of reads aligned to a concatenated dual reference genome or C. difficile genome of all sequenced samples.**

| Sample type | Number of reads | Aligned to concatenated genome reference (%) | Aligned to *C. difficile* genome (%) |
| --- | --- | --- | --- |
| Bacterial control 1 | 8936289 | 99.11 | 99.21 |
| Bacterial control 2 | 12094137 | 99.02 | 99.40 |
| Bacterial control 3 | 70755533 | 99.33 | 99.55 |
| Human control 3 h 1 | 19115353 | 94.38 | 0.00 |
| Human control 3 h 2 | 25470128 | 90.68 | 0.00 |
| Human control 3 h 3 | 27025069 | 95.67 | 0.00 |
| Human control 6 h 1 | 21468597 | 91.70 | 0.00 |
| Human control 6 h 2 | 24593073 | 92.11 | 0.00 |
| Human control 6 h 3 | 57975499 | 96.20 | 0.00 |
| Human control 12 h 1 | 22390842 | 91.97 | 0.00 |
| Human control 12 h 2 | 23882910 | 94.54 | 0.00 |
| Human control 12 h 3 | 58915160 | 96.13 | 0.00 |
| Human control 24 h 1 | 23835563 | 90.63 | 0.00 |
| Human control 24 h 2 | 21564602 | 93.29 | 0.00 |
| Human control 24 h 3 | 45215959 | 94.42 | 0.00 |
| 3 h infected 1 | 49226860 | 95.45 | 1.32 |
| 3 h infected 2 | 58246564 | 97.60 | 0.19 |
| 3 h infected 3 | 55316199 | 96.99 | 1.68 |
| 6 h infected 1 | 57700752 | 97.21 | 0.14 |
| 6 h infected 2 | 58987014 | 98.14 | 0.06 |
| 6 h infected 3 | 57515435 | 97.75 | 0.32 |
| 12 h infected 1 | 56507425 | 97.25 | 0.33 |
| 12 h infected 2 | 53660909 | 98.25 | 0.11 |
| 12 h infected 3 | 53241414 | 97.93 | 0.42 |
| 24 h infected 1 | 62622140 | 97.76 | 2.06 |
| 24 h infected 2 | 64673740 | 97.50 | 0.06 |
| 24 h infected 3 | 54650378 | 96.98 | 0.44 |
